# Freshwater dissolved oxygen dynamics: changes due to glyphosate, 2,4-D and their mixture, both under clear and turbid-organic conditions

**DOI:** 10.1101/2021.09.02.458549

**Authors:** V.L. Lozano, C.E. Miranda, A.L. Vinocur, C.A. Sabio y García, M.S. Vera, C. González, M.J. Wolansky, H.N. Pizarro

**Author notes:** FCEN, UBA. Int. Güiraldes 2160, Pabellon II, C1428EHA, Buenos Aires, Argentina.

## Abstract

We performed two independent outdoor mesocosm experiments where we measured the variation of DO saturation (DO%) in freshwater after a single input of Roundup Max^®^ (G) (glyphosate-based formulation), AsiMax 50^®^ (2,4-D) (2,4-D-based formulation) and their mixture (M). Two concentration levels were tested; 0.3 mg/L G and 0.135 mg/L 2,4-D (Low; L) and 3 mg/L G and 1.35 mg/L 2,4-D (High; H). We assayed consolidated microbial communities coming from a system in organic turbid eutrophic status and a system in clear mesotrophic status during 21 and 23 days, respectively. A sample of phytoplankton (micro+nano, pico-eukaryotes, pico-cyanobacteria), mixotrophic algae and heterotrophic bacteria was collected to determine abundances at each of four sampling dates. The clear and turbid systems showed similar, but not synchronized, patterns of daily DO% changes in relation to the controls (DO%_v_), after exposure to both single and combined formulations. Under glyphosate scenarios (GL, GH, ML and MH), the two types of systems showed similar DO%_v_ but different microbial abundances, being associated to an increase in the micro+nano and pico-eukaryotic phytoplankton fractions for the clear system. In contrast, in the turbid system changes were associated with increased pico-cyanobacteria and decreased mixotrophic algae. Effects of 2,4-D were only observed in the turbid system, leading to decreased micro+nano phytoplankton abundances. Under the turbid scenario, the herbicide mixture at high concentration had a synergistic effect on DO%_v_ and recovery was not detected by the end of the experiment. Our results revealed that herbicides inputs induced changes in phytoplankton abundances that leads to measurable DO variations.

## 1. Introduction

In aquatic systems, dissolved oxygen (DO) integrates the dynamics of physical, chemical and biological variables, and its concentration results from the balance of several processes. With regard to biological processes, DO results from the balance between photosynthesis and respiration. Photosynthesis by autotrophic freshwater communities (eg., phytoplankton, attached microbial communities and macrophytes) is a major source of oxygen for all living organisms, while oxygen is consumed by the respiration of free and attached bacteria, phyto and zooplankton, attached auto- and heterotrophic communities, macroinvertebrates and vertebrates. Moreover, DO concentration in freshwater systems may be affected by biogeochemical processes such as mineralization of allochthonous organic matter and nitrification. For these reasons, DO has been fully considered as an ecological indicator.

Therefore, DO monitoring in natural freshwater systems is a useful tool for different applications, including management, conservation or restoration.

The techniques for measuring DO have evolved from chemical reactions (e.g., Winkler method) to the use of microelectrodes and optical sensors. The latter are small, cheap and have long-term stability (Klimant et al., 1995). Currently, monitoring of DO in aquatic systems, both *in situ* and under laboratory conditions, is easy providing instant results (Warkentin et al., 2007).

In Argentina, about 100,000 shallow lakes are located in the Pampa plain, one of the largest and most productive agroecosystems worldwide (Dukatz et al., 2006). Most of these shallow lakes alternate between two contrasting limnological states: the clear state, where the primary productivity is mainly driven by submerged macrophytes; and the turbid state, which shows prevalence of phytoplanktonic productivity (Allende et al., 2009). Some studies have provided evidence that shallow lakes remain turbid due to agrochemical contamination, caused by fertilizers (Quirós et al., 2002) and herbicides such as glyphosate (Vera et al., 2010, Castro Berman et al., 2018).

There was an increase in agrochemical-based agriculture between 1995 and 2015 worldwide, which was accompanied by an increase in the use of insecticides and herbicides of about 3.5 and 4.4 times, respectively (FAOSTAT, 2017). Scientific results show that agrochemicals exert direct and indirect toxic effects on freshwater organisms living in agricultural landscapes (Lampert et al., 1989; Fleeger et al., 2003). The ecosystem approach is broadly accepted as the most realistic framework to investigate the impact of anthropogenic disturbances on freshwater organisms, while DO has historically been used as a reliable indicator of water quality. In this regard, King et al. (2015) pointed out that DO is particularly sensitive to herbicide contamination in water.

In Argentina, glyphosate (N-phosphonomethylglycine) and 2,4-D (2,4-dichlorophenoxyacetic acid) are the most widely used herbicides. They show different modes of action. Glyphosate is a reversible inhibitor of EPSP (5-enolpyruvylshikimate-3-phosphate) synthase, a key enzyme in the synthesis of aromatic amino acids in plants, algae, bacteria and fungi (Pollegioni et al., 2011). On the side, 2,4-D is an auxin-type herbicide, which causes the death of dicotyledonous by inducing overgrowth of vascular cambium (Song, 2014). The appearance of glyphosate-resistant weeds due to its continued extensive use prompted farmers to apply mixtures as a control option. An experimental laboratory study reported that the mixture of these herbicides has a synergistic effect on freshwater cyanobacteria (Lozano et al., 2018). However, the environmental implications of their combined application have been poorly investigated.

The aim of the present research was to study the daily changes in dissolved oxygen saturation percentage (DO%) resulting from the impact of Roundup Max^®^, (a glyphosate-based herbicide formulation) and AsiMax 50^®^ (a 2,4-D-based herbicide formulation), alone and in mixture, on two freshwater systems of different turbidity. We carried out two outdoor mesocosm experiments to analyze the daily DO% variation with respect to control (DO%v) in water, and the difference in DO%_v_ between a single dose of Roundup Max^®^, AsiMax 50^®^ or their mixture with respect to the controls, under different turbidity conditions, i.e. clear and organic turbid. High and low concentrations of each herbicide and their respective mixtures were assayed to assess a hypothetical dose-dependence effect. A sample was obtained on each of four occasions during the experiment to determine the abundance of the following communities: micro+nano phytoplankton (>2 µm, M+N); photosynthetic picoplankton (PPP, 0.2-2 µm), which includes prokaryotic picocyanobacteria (Pcy) and eukaryotic phototrophs (PEuk); and heterotrophic bacterioplankton (HB).

Algae and cyanobacteria are possibly sensitive to herbicide contamination because they share several metabolic pathways with vascular plants (Ferrari et al., 2018). In addition, we may assume that glyphosate and 2,4-D have different mode of action but similar toxicological endpoint. Then, it is reasonable to predict that their combination will show additive effects, as postulated by Faust et al. (2001). On the other hand, photosynthetic microbial communities are more abundant in organic turbid than in clear systems. Based on these considerations, we tested the following hypotheses:

1. Daily DO%_v_ pattern are similar for the organic turbid system and for the clear one after exposure to Roundup Max^®^ and AsiMax 50^®^, alone and in mixture.
2. The impact of Roundup Max^®^ and AsiMax 50^®^ on the daily pattern of DO%_v_ are similar.
3. The effect of the mixture of Roundup Max^®^ and AsiMax 50^®^ on the daily pattern of DO%_v_ is similar to the sum of the effects of the herbicides acting independently (additive effect).
4. The impact of the herbicides on the dynamics of daily DO%_v_ is higher at high doses than at low doses.
5. DO%_v_ pattern resulting from the impact of Roundup Max^®^, AsiMax 50^®^ and their mixture may be explained by phytoplanktonic abundance variations.

## 2. Material and methods

### 2.1. Experimental design

Natural freshwater systems in the Pampean plain of Argentina are exposed to an intensive herbicide spraying and the possibility of contamination cannot be ruled out. Therefore, to obtain microbial communities without previous exposition, we established freshwater systems through colonization in outdoor tanks of 3,000 L, allowing them to evolve for two years. This decision was taken for avoiding the use of natural lake water where possible previous contamination is impossible to be guaranteed. We performed two outdoor mesocosm experiments using these consolidated microbial communities coming from systems developed in two tanks: (a) a system in organic, turbid eutrophic status (hereafter referred to as turbid) with chlorophyll *a* concentration = 31.7 µg/L and turbidity = 12 NTU; and; (b) a system in clear mesotrophic status (hereafter referred to as clear) with chlorophyll *a* concentration = 8.7 µg/L and turbidity = 2 NTU. The turbid experiment was carried out 21 d in late spring, while the clear one was performed 23 d in late summer, using a similar experimental design. Experimental units (EUs) were 40-45-L plastic bags incubated in three 3,000-L outdoor tanks located in the experimental campus at the University of Buenos Aires. The EUs of the first experiment were filled with turbid water and the EUs of the second one with clear water, both of which were taken from the natural colonized systems. Prior to the assays, the EUs were subjected to a 3-day stabilization period and randomly assigned to one of the following treatments: Control (C); a single dose of Roundup Max^®^ (glyphosate-based formulation, active ingredient, a.i.: G) at low (0.3 mg/L G) and high (3 mg/L G) concentrations; a single dose of AsiMax 50^®^ (2,4-D-based formulation, a.i.: 2,4-D) at low (0.135 mg/L 2,4-D) and high (1.35 mg/L 2,4-D) concentrations; and a herbicide mixture at two concentrations with the same ratio of active ingredients (i.e. 1 G : 0.45 2,4-D), namely a low mixture concentration of 0.3 mg/L G + 0.135 mg/L 2,4-D and a high mixture concentration of 3 mg/L G + 1.35 mg/L 2,4-D). Hereafter, these treatments will be referred to as Control, GL, GH, 2,4-DL, 2,4-DH, ML and MH, respectively. Treatments were realized by simple overthrow andperformed in triplicate, except for Control, which was replicated six times (two per tank). The summary of the experimental design is shown in Fig. 1.

**Fig. 1:**
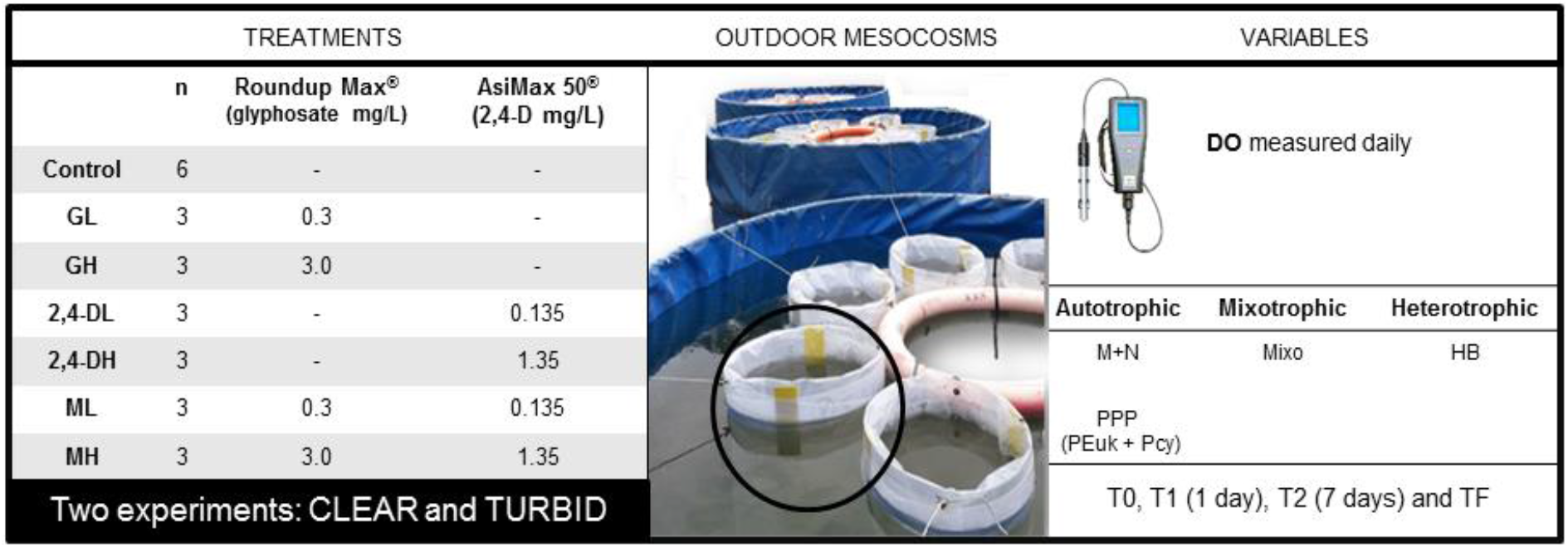
Scheme of the experimental design of clear and turbid assays. Treatments are listed on the left and variables on the right. GL: glyphosate low concentration, GH: glyphosate high concentration, 2,4-DL: 2,4-D low concentration, 2,4-DH: 2,4-D high concentration, ML: mixture low concentration, MH: mixture high concentration, M+N: micro+nano phytoplankton, PPP: photosynthetic picoplankton, PEuk: eukaryotic phototrophs, Pcy: prokaryotic picocyanobacteria, Mixo: mixotrophic micro+nano plankton, HB: heterotrophic bacteria. Black circle: mesocosm.

### 2.2. Variables measured in clear and turbid experiments

#### 2.2.1. Environmental variables

Daily maximum and minimum data of air temperature were determined from the National Weather Service of Argentina (Servicio Metereológico Nacional, SMN), using available data from a station at less than 2 km away from the experimental campus. Values were used to calculate daily mean air temperature (T°C) defined as (T_maximum_ + T_minimum_)/2. An *ad-hoc* device of the same shape and dimensions as those of an EU was placed near the tanks during the whole study periods to measure rainfall volume (in mL).

#### 2.2.2. Chemical variables

The concentration of DO and DO% was monitored daily at 8:30 AM with an optical dissolved oxygen meter (YSI^®^ ProODO™; error = 0.01 mg/L) by immersing the probe to a depth of 10 cm in the center of each EU. Water temperature was continuously monitored using thermo buttons (Akribis^®^) immersed in each EU. For herbicides analysis, 50 mL of water was taken from the center of each EU with plastic flasks and kept at -20°C. Glyphosate was determined by ELISA Microtiter Plate test (Abraxis^®^) and 2,4-D by HPLC-UV (APHA, 2005; U.S.EPA, 1980) on days 0, 1, 7 and 21 in the turbid experiment and on days 0, 1, 7 and 23 in the clear experiment. Half-lives were calculated considering that glyphosate and 2,4-D residue adjusted to a logarithmic function assuming a first-order kinetic.

#### 2.2.3. Biological variables

##### 2.2.3.1. M+N phytoplankton abundances

Total abundances of live micro (>20 µm) and nano (2–20 µm) phytoplankton (M+N) and mixotrophic micro + nanophytoplankton (Mixo) fractions were analyzed with inverted microscope. For this purpose, 200 mL of water was sampled from each EU at T0 (immediately after treatment application), T1 (1 d), T2 (7 d) and TF (21 and 23 d). Samples were fixed with 1% acidified Lugol’s iodine solution for algal and cyanobacteria quantification (> 2 µm) following the technique of Utermöhl (1958). In general, Bourrelly’s systematic scheme (1970) was followed for taxonomic identification. *Peridinium* sp. and *Ochromonas* sp. were considered to be mixotrophic organisms (Andersson et al., 1989, Palsson & Granéli, 2004)

##### 2.2.3.2. PPP and HB abundances

Total abundances of photosynthetic picoplankton (PPP) (0.2–2 µm), which includes picocyanobacteria (Pcy) and eukaryotic phototrophs (PEuk), and heterotrophic bacteria (HB), were obtained by flow cytometry. A FACSAria II (Becton Dickinson^®^) flow cytometer equipped with a standard 15 mW blue argon-ion (488 nm emission) laser and a red laser diode (635 nm) was used. Samples (3.6 mL) were taken on the same sampling dates as the M+N and Mixo samples. They were fixed with 10% cold glutaraldehyde (1% final concentration), left in the dark for 10 min at room temperature and then stored at -80°C. A known volume of beads (1-µm diameter, Fluospheres^®^) was added to all unfrozen samples. PPP were identified in plots of SSC (Side Scatter) versus blue laser-dependent red fluorescence (PerCP or FL3, 670 nm), orange fluorescence (PE or FL2 585/42 nm) versus FL3, and red laser-dependent far-red fluorescence (APC or FL4, 661 nm) versus FL3. For the HB determination, 400 µL-samples were previously stained with SYBRGreen I (Sigma-Aldrich^®^) diluted in DMSO to a final concentration of 1X, left for about 10 min in the dark to complete the nucleic-acid staining and run in the flow cytometer. Abundances were calculated by their signature in plots of side scatter light (SSC) versus green fluorescence of nucleic acid-bound stains FITC (FL1 530 nm) and SSC (side scatter) axes (Gasol et al. 1999).

### 2.3. Statistical and numerical analysis

A *t-*test was used to compare between the real values of G and 2,4-D for each treatment at T0.

Given that DO depends on temperature and that both experiments were carried out at different times of the year, we evaluated the effect of herbicides on the daily DO percentage of saturation variation (DO%_v_), calculated as DO%_v_ = DO%_x_ – DO%_control_, where DO%_x_ is the value obtained from each treatment with herbicide (x). Then, DO%_v_ can attain positive or negative values. Differences in DO%_v_ among the seven treatments over time were analyzed with one-way RM-ANOVA with *treatment* as the factor with 7 levels (GL, 2,4-DL, ML, GH, 2,4-DH, MH and C) and 21 (turbid experiment) and 23 (clear experiment) measurement days. Statistical significance was set at p<0.05, considering periods of at least 3 days showing significant differences compared with the controls. The three tanks served as replicates.

A one-way RM-ANOVA was performed to test possible additive effects on DO%_v_, with *mixture* as the factor with 2 levels: observed and expected and 21 (turbid experiment) and 23 (clear experiment) measurement days, for both clear and turbid experiments. Expected values were obtained by summing the values of DO%_v_ measured for each herbicide separately. When the difference between the observed and expected DO%_v_ was significant, the interaction was synergistic if the observed mean DO%v concentration was higher than the expected mean DO%_v_ concentration; otherwise, the interaction was antagonistic (Lozano et al., 2018).

A one-way RM-ANOVA was conducted for M+N, Mixo, as well as for PPP and HB abundances with *treatment* as the factor with 7 levels (GL, 2,4-DL, ML, GH, 2,4-DH, MH and C) and 21 (turbid experiment) and 23 (clear experiment) sampling days. All data were previously analyzed for normality and homocedasticity using the Shapiro-Wilks and Levene tests, respectively, and log-transformed if appropriate.

Pearson correlation matrix was calculated among the abundance values of the autotrophic, mixotrophic and heterotrophic fractions and DO%, for both experiments. With these correlation values, graphical network was built in Cytoscape^®^ v3.7.1 (Shannon et al. 2007). Thickness of lines was set proportionally to the r values obtained while the size of the circles was set proportionally to the mean values of abundance of the different biological fractions from the controls (ind/mL) in each system at the initial time.

Statistical analyses were performed using SigmaPlot^®^ 11.0 and InfoStat^®^ software version 2015.

## 3. Results

### 3.1. Environmental variables

As expected for independent experiments made in different moments, there were differences in climatic conditions throughout the study periods, which are shown in Fig. 2. Daily mean air temperature was 21.9°C (min. 15.0 and max. 24.1°C) for the clear experiment and 17.9°C (min. 13.8 and max. 23.2°C) for the turbid one. Rainfall volume in each EU was 7,840 mL (6 rain episodes) for the entire clear experiment and 7,275 mL (6 rain episodes) for the turbid one.

**Fig. 2:**
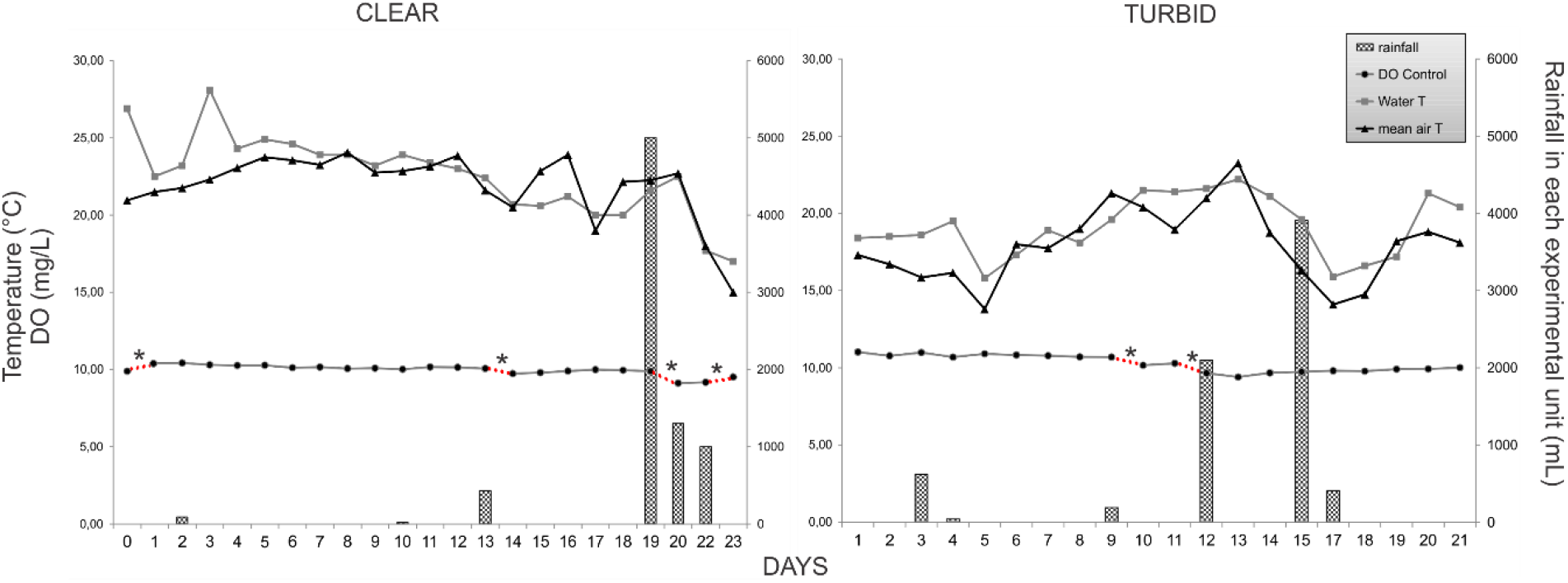
Mean daily DO concentration calculated from control EUs (n=6) and mean air and water temperature (°C) values and rainfall (mL) accumulated in each EU for the clear and turbid experiments along the study period. Significant differences in DO values (RM-ANOVA, p<0.05) are shown in dotted red lines with asterisk (*).

### 3.2. Physical and chemical variables

Initial herbicide concentrations were similar to the nominal concentration applied (Table 1); no differences in herbicide concentrations were found between clear and turbid experiments for all treatments at initial time (*t-*test, p>0.1 in all cases). The half-lives of herbicides where 16 and 41 days for glyphosate and 99 and 69 for 2,4-D in the clear and turbid experiments respectively. Water temperature measured in EUs using thermos-buttons showed a mean daily variation between maximum and minimum values of 2.32 ± 0.68 °C in each EU for the clear experiment, and 2.55 ± 0.92 °C for the turbid experiment.

**Table 1:** Comparison between nominal and mean real values (± 1 SD) of herbicide lives for both herbicides during the two experiments. LOD: Limit of detection.

| Nominal (mg/L) |  |  | Real (mg/L) |  |  |  |
| --- | --- | --- | --- | --- | --- | --- |
|  |  |  | CLEAR |  | TURBID |  |
|  | G | 2,4-D | G | 2,4-D | G | 2,4-D |
| C | 0 | 0 | < LOD | < LOD | < LOD | < LOD |
| GL | 0.30 | 0 | 0.28 ± 0.06 | < LOD | 0.36 ± 0.08 | < LOD |
| 2,4-DL | 0 | 0.135 | < LOD | 0.11 ± 0.01 | < LOD | 0.14 ± 0.01 |
| ML | 0.30 | 0.135 | 0.25 ± 0.05 | 0.12 ± 0.01 | 0.33 ± 0.05 | 0.11 ± 0.01 |
| GH | 3 | 0 | 3.09 ± 0.08 | < LOD | 3.14 ± 0.09 | < LOD |
| 2,4-DH | 0 | 1.35 | < LOD | 1.02 ± 0.07 | < LOD | 1.28 ± 0.02 |
| MH | 3 | 1.35 | 2.93 ± 0.33 | 1.05 ± 0.13 | 2.82 ± 0.43 | 1.26 ± 0.08 |
| Half-lives (days) |  |  | 16 | 99 | 41 | 69 |

Maximum and minimum absolute values of DO concentrations (mg/L) in EUs treated with herbicides ranged between 8.15 mg/L (MH, day 20) and 11.00 mg/L (GH, day 8) for the clear system, and between 8.52 mg/L (MH, day 13) and 12.09 mg/L (GH, day 6) for the turbid one. The percentage of oxygen saturation always exceeded 94% in both water systems. In regard to daily DO concentrations in the controls, the clear system appeared to be more affected by rainfall periods than the turbid one (Fig. 2). The maximum DO%_v_ was observed in the clear system with a mean increase of +10.30% under the GH treatment, while the minimum percentage was detected in the turbid system with a mean decrease of -9.17% under the MH treatment. For G, the turbid system showed a slight but significant increase in mean DO%_v_ of +2.60% (±0.44%) at 19, 20 and 21 d under the GL treatment (p=0.012, p=0.002 and p<0.001, respectively), while no increase was observed for the clear system (Fig. 3a and 4a). In the GH treatment under turbid experiment, DO%_v_ increased between days 3 and 9 with a mean of +6.56% (±1.35%), and between 14 and 21 d with a mean of +3.59% (±0.69%) (p<0.001 in all cases, Fig. 4b); a similar increase of +6.33% (±3,10%), was recorded a few days later, between days 7 and 12, under clear conditions (Fig. 3b). The maximum percentage of DO%_v_ was reached under the GH treatment, with an average increase of +7.92% at day 6 (p<0.001) for the turbid and +10.30% at day 8 (p<0.001) for the clear experiments. Although DO%_v_ increased similarly in both systems, the impact started earlier and lasted longer under the turbid condition.

**Fig. 3:**
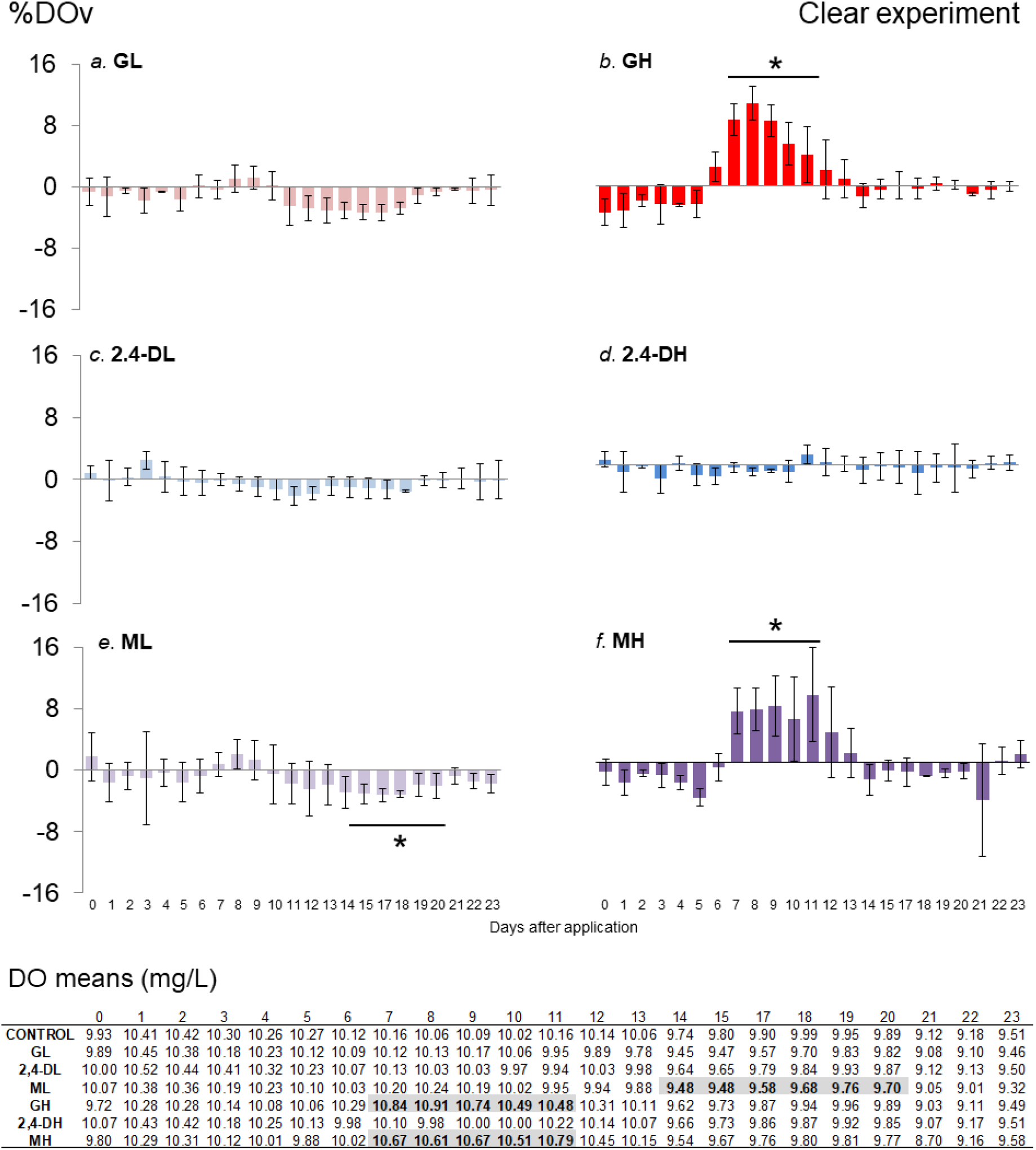
Daily percentage of DO saturation in relation to the controls (DO%_v_) in the clear experiment after herbicide application (G: Roundup Max^®^; 2,4-D: AsiMax 50^®^; M: Roundup Max^®^ + AsiMax 50^®^; L: low concentration; H: high concentration). Asterisks indicate significant differences with controls (RM-ANOVA, p<0.05). Below: Daily DO mean values (mg/L) for each treatment; in grey and bold: values corresponding to significant differences showed in the graphs below (RM-ANOVA, p<0.05).

**Fig. 4:**
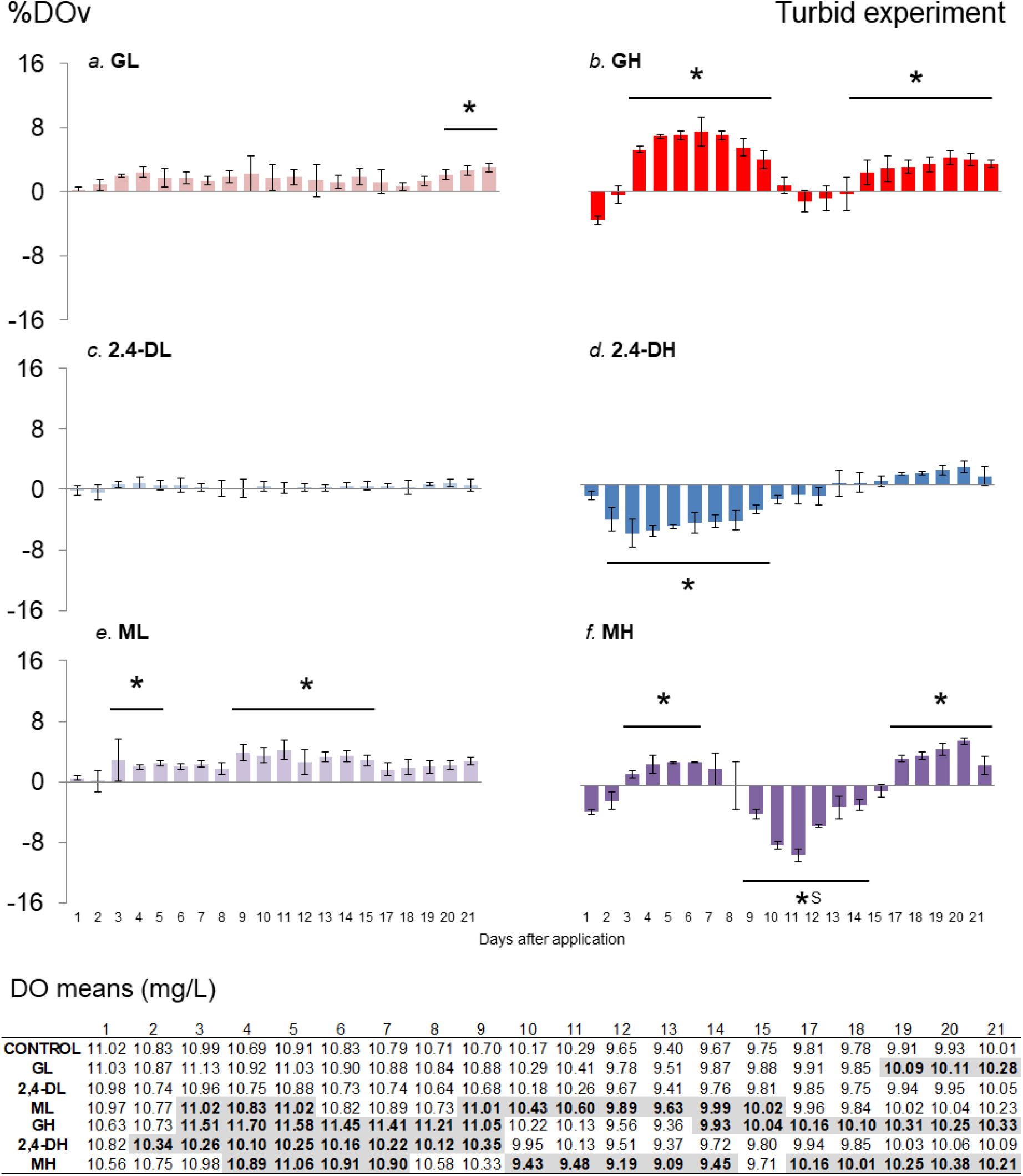
Daily percentage of DO saturation in relation to the controls (DO%_v_) in the turbid experiment after herbicide application (G: Roundup Max^®^; 2,4-D: AsiMax 50^®^; M: Roundup Max^®^ + AsiMax 50^®^; L: low concentration; H: high concentration). Asterisks indicate significant differences with controls (RM-ANOVA, p<0.05) and (S): synergism. DO means values (mg/L) are listed below, in grey and bold: values corresponding to significant differences showed in the graphs below ((RM-ANOVA, p<0.05).

The addition of 2,4-D to the water did not modify DO%_v_ in all treatments, except for a decrease observed under the 2,4-DH treatment in the turbid system (p<0.001 for all days), observed between days 2 and 9 with a mean of -5.14% (±0.98%) (Fig. 4d).

The mixture of herbicides at low and high doses impacted DO%, under both clear and turbid conditions. Under turbid conditions, DO%_v_ increased in the ML treatment 3, 4 and 5 days after application with a mean of +2.44% (±0.46%) (p<0.001, p=0.02 and p=0.004, respectively) and between days 9 and 15 with a mean value of +3.39 (±0.56%) (p<0.001 for all days except day 12 with p=0.002) (Fig. 4e). Under clear conditions, ML treatment reduced DO in the period of 14 to 20 days (Fig. 3e). In the clear system, MH increased DO%_v_ by +6.19% (±1.60%) between days 7 and 12 (p<0.001 for all days, except day 12 with p=0.005) (Fig. 3f). In the turbid system, MH increased DO%_v_ by +2.83% (±0.45%) at days 4-6 (p<0.001) and 7 (p=0.005), followed by a decrease of -5.22% (±2.75%) between days 9 and 14. Thereafter, DO%_v_ incresed by +4.38% (±1.35%) from day 18 onwards (p<0.001 for all days) (Fig. 4f).

At the end of the experiment, the herbicide treatments under the clear condition had DO% values similar to those of the controls, suggesting a recovery of the system. A contrasting result was obtained for the turbid system, where the treatments with glyphosate (GL, GH and MH) showed a net significant increase in DO%_v_ (Fig. 4a,b,f). Compared with the other glyphosate-related treatments, the MH treatment in turbid experiment trended toward no recovery to control levels of DO%_v_ at the end of the study (Fig. 4e).

In general, the combined exposure to G and 2,4-D formulations caused additive effects on the examined variables, except for MH in the turbid system, which exhibited significant synergism (p<0.05) between days 9 and 14 (Fig. 4f).

### 3.3. Biological variables

In the clear experiment, the mean abundance of M+N ranged from a minimum of 3.4 × 10^2^ ± 3.1 × 10^2^ ind/mL (GL, TF) to a maximum of 1.8 × 10^4^ ± 6.8 × 10^3^ ind/mL (GH, T2). This microbial fraction showed behavioral differences among treatments over the study period. In general, there was a trend toward a significant increase in abundance under treatments GL, ML, GH and MH. On the contrary, M+N showed a trend toward decreased abundance under treatments Control, 2,4-DL and 2,4-DH, probably as a consequence of an enclosure effect. The abundance of PEuk ranged from a minimum of 4.2 × 10^2^ ± 3.4 × 10^2^ ind/mL (2,4-DL, TF) to a maximum of 1.5 × 10^4^ ± 7.8 × 10^3^ ind/mL (GH, T2). For all treatments, this microbial fraction showed similar abundance values between TF and T0, with intermediate patterns. The abundance of Pcy ranged from a minimum of 4.6 × 10^4^ ± 1.3 × 10^4^ ind/mL (GH, T0) to a maximum of 2.4 × 10^5^ ± 1.7 × 10^5^ ind/mL (GH, TF). Likewise, the abundance values of Pcy were similar between Tf and T0 for all treatments. The abundance of Mixo ranged from a minimum of 1.4 × 10^1^ ± 2.3 × 10^1^ ind/mL (GH, TF) to a maximum of 7.1 × 10^3^ ± 3.4 × 10^3^ ind/mL (2,4-DL, T0). The abundances of this algal fraction showed a decreasing trend toward TF for all treatments, except for ML and MH where abundance values were similar between T0 and TF. The abundance of heterotrophic bacteria ranged between a minimum of 6.1 × 10^6^ ± 1.3 × 10^6^ ind/mL (Control, T1) and a maximum of 2.6 × 10^7^ ± 3.6 × 10^6^ ind/mL (Control, T2) (Supplementary Material 1)

When compared with controls, significant treatment-related effects on M+N were observed at T2, for both GH with a mean of +769% (ANOVA, p<0.001) and MH with a mean of +267% (ANOVA, p=0.01) (Fig. 4a). These increases persisted until TF, with +125% for MH (ANOVA, p<0.001) and +55% for GH (ANOVA; p=0.004). At TF, GL showed a decrease of -83% in relation to the controls (ANOVA, p=0.001). For PEuk, abundance increased at T2 in GH treatment with a mean value of 414% and in MH with an increase of 204%. No treatment-related effects were observed for Pcy and Mixo fractions. HB fraction had consistent variations in abundance across treatments at various sampling times: a mean increase of +52% in GH (p<0.001) at T0; +60, +49 and +136% (p<0.001 in all cases) in 2,4-DH, GH and MH, respectively at T1. In contrast, some treatments showed lower abundances than controls at T2, with -26% (p=0.011), -27% (p=0.008) and -27% (p=0.009) for GH, 2-4-DH and MH, respectively (Fig. 5e).

**Fig. 5:**
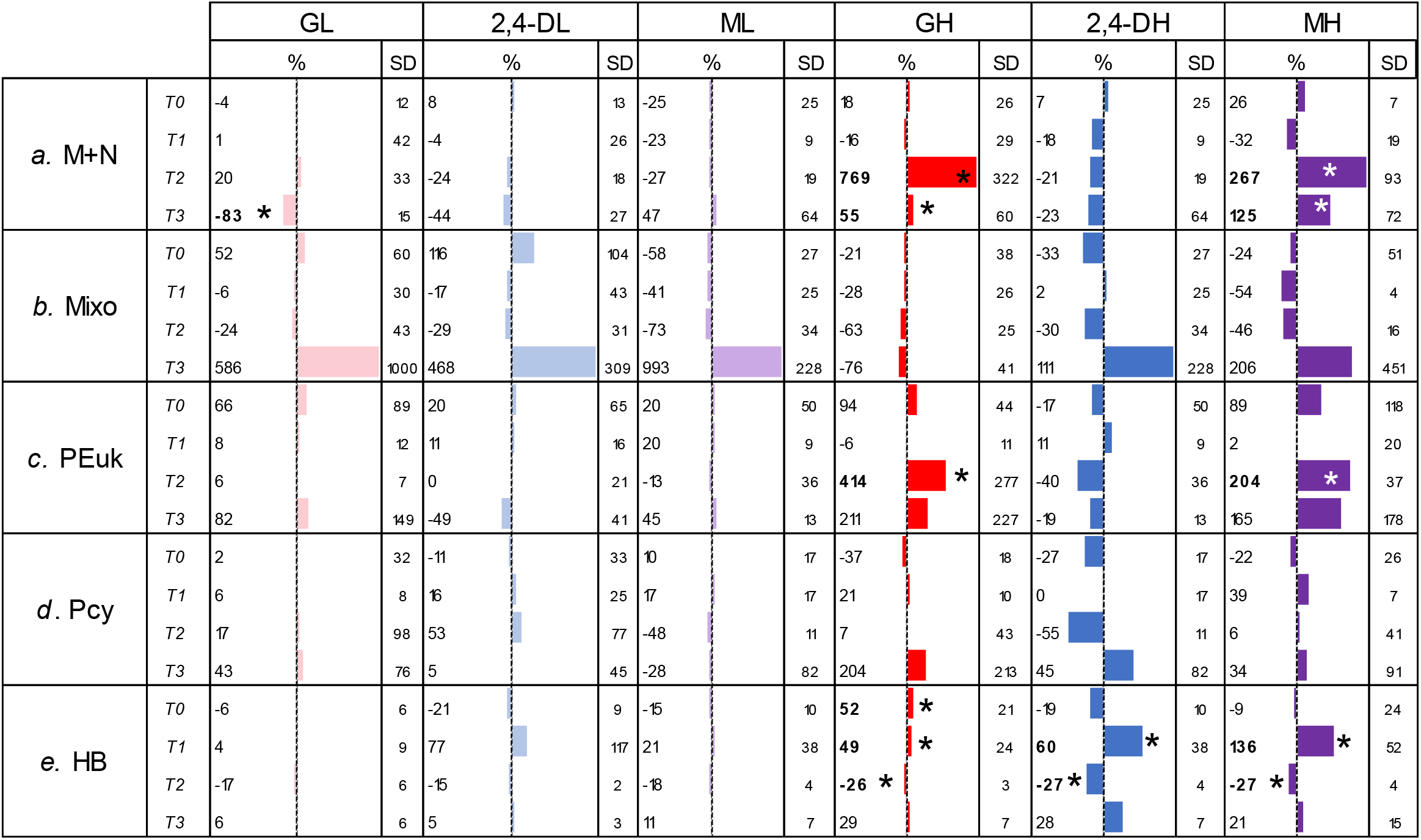
Percentages (%) and standard deviation (SD) of the abundance of microbial communities for each treatment in the clear system. M+N: micro+nano phytoplankton; PEuk: picoeukaryotes; Pcy: picocyanobacteria; Mixo: mixotrophic algae and HB: heterotrophic bacteria. (G: Roundup Max^®^; 2,4-D: AsiMax 50^®^; M: Roundup Max^®^ + AsiMax 50^®^; L: low concentration; H: high concentration). Bars show the mean % (n=3) around controls (central line). Asterisks indicate significant differences with respect to controls for each microbial fraction (RM-ANOVA, p<0.05).

In the turbid experiment the mean abundances of M+N ranged from a minimum of 1.1 × 10^4^ ± 3.5 × 10^3^ ind/mL (2,4-DH, TF) to a maximum of 1.6 × 10^5^ ± 4.9 × 10^4^ ind/mL (GL, T2). There was a trend toward decreased abundance under treatments Control, GL, ML and 2,4-DH, where values at TF were lower than those at T0. The abundance of PEuk varied from a minimum of 3.3 × 10^4^ ± 5.0 × 10^3^ ind/mL (GL, TF) to a maximum of 2.3 × 10^5^ ± 7.2 × 10^3^ ind/mL (2,4-DH, T3). The abundance of the Pcy fraction varied from a minimum of 2.0 × 10^5^ ± 3.9 × 10^4^ ind/mL (GL, TF) to a maximum of 8.2 × 10^5^ ± 2.7 × 10^5^ ind/mL (MH, T3). Mixo abundance ranged from a minimum of 3.0 × 10^3^ ± 3.2 × 10^3^ ind/mL (MH, TF) to a maximum of 5.5 × 10^4^ ± 2.2 × 10^4^ ind/mL (Control, T3). The abundance of heterotrophic bacteria ranged between a minimum of 2.5 × 10^6^ ± 6.4 × 10^5^ ind/mL (Control, T0) and a maximum of 1.4 × 10^7^ ± 2.6 × 10^6^ ind/mL (ML, T3) (Supplementary Material 2).

In relation to controls, M+N showed significant decreases in 2,4-DH both at T2 and TF, with mean values of -67% (ANOVA, p<0.001) and -67% (ANOVA, p=0.003), respectively (Fig. 6a). The abundance of Mixo showed significant decreases with respect to control at T2 in GH (−73%), MH (−72%), 2,4-DH (−64%), ML (−50%) and 2,4-DL (−49%) (ANOVA, p<0.001 in all cases). Only MH showed a sustained decrease in abundance of -88% toward TF (ANOVA, p=0.006) (Fig. 6b). PEuk abundance values showed significant treatment-related effects for GH at T1, with a mean variation of -42% (ANOVA, p=0.007), and for GL, 2,4-DH and MH at TF, with mean variations of -60% (ANOVA, p >0.001), +58% (ANOVA, p=0.004) and +53% (ANOVA, p=0.004) respectively (Fig. 6c). Pcy abundance in ML and GH increased slightly with respect to Control, with means of +33 and +34% (ANOVA, p=0.005 and 0.007, respectively) at T1. There were significant increases in Pcy abundance in GH and MH from T2 to TF, with means of +58 and +104% for GH, and +89 and +98% for MH (ANOVA, p<0.001 in all cases). In addition, ML showed an increase in the abundance of the Pcy fraction at T1 and TF, with a mean of +33 and +55% respectively (ANOVA, both p<0.001) (Fig. 6d). HB showed rapid variations with respect to control at T0 and, except for GL, all treatments showed increases until TF: 2,4-DL +67%, ML +90%, GH +157%, 2,4-DH +94% and MH 250% (ANOVA, p<0.001 in all cases) (Fig. 6e).

**Fig. 6:**
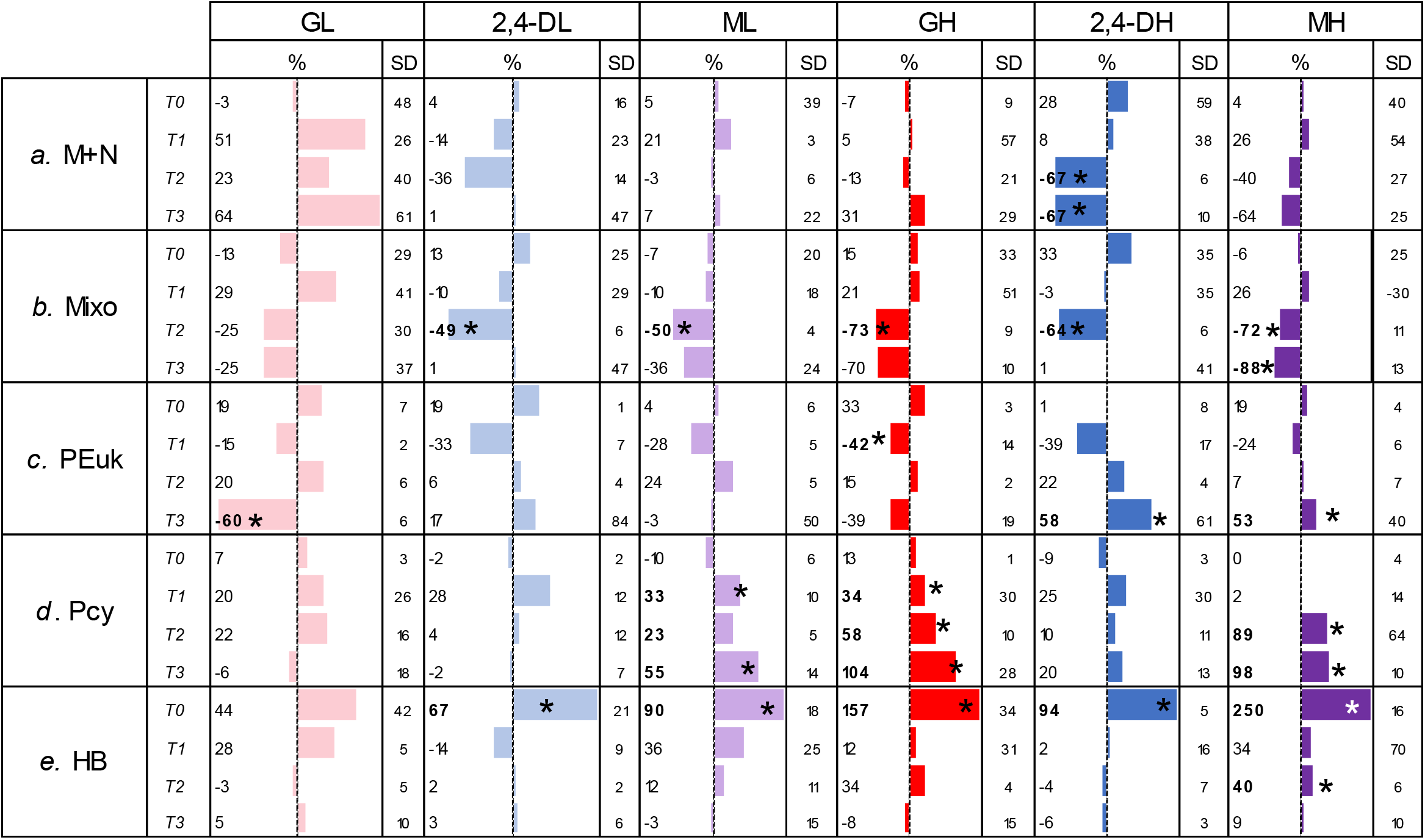
Percentages (%) and standard deviation (SD) of the abundance of microbial communities for each treatment in the clear system. M+N: micro+nano phytoplankton; PEuk: picoeukaryotes; Pcy: picocyanobacteria; Mixo: mixotrophic algae and HB: heterotrophic bacteria. (G: Roundup Max^®^; 2,4-D: AsiMax 50^®^; M: Roundup Max^®^ + AsiMax 50^®^; L: low concentration; H: high concentration). Bars show the mean % (n=3) around controls (central line). Asterisks indicate significant differences with respect to controls for each microbial fraction (RM-ANOVA, p<0.05).

To analyze the relationship between DO% and variations in the abundance of all the fractions of autotrophic and bacterial communities, a correlation network was built with these variables at T0, T1, T2 and TF by means of Cytoscape^®^ using Pearson’s r values (Fig. 7). Under clear conditions, DO% correlates positively with M+N phytoplankton abundances (r=0.53, p<0.0001), which, in turn correlates positively with PEuk abundances (r=0.54, p=0.0001). On the other hand, HB correlates negatively with DO% (r=-0.24, p=0.0408). Under turbid conditions,, DO% correlates positively with Pcy abundances (r=0.38, p=0.0044) and negatively with mixotrophic algae (r=-0.36, p=0.008). In regard to relationships between abundance variations, Pcy correlates negatively with Mixo (r=-0.52, p<0.0001), which also correlates negatively with M+N phytoplankton (r=-0.5, p=0.0001). Finally, mixotrophic algae correlates positively with M+N phytoplankton (r=0.43, p= 0.0001).

**Fig. 7:**
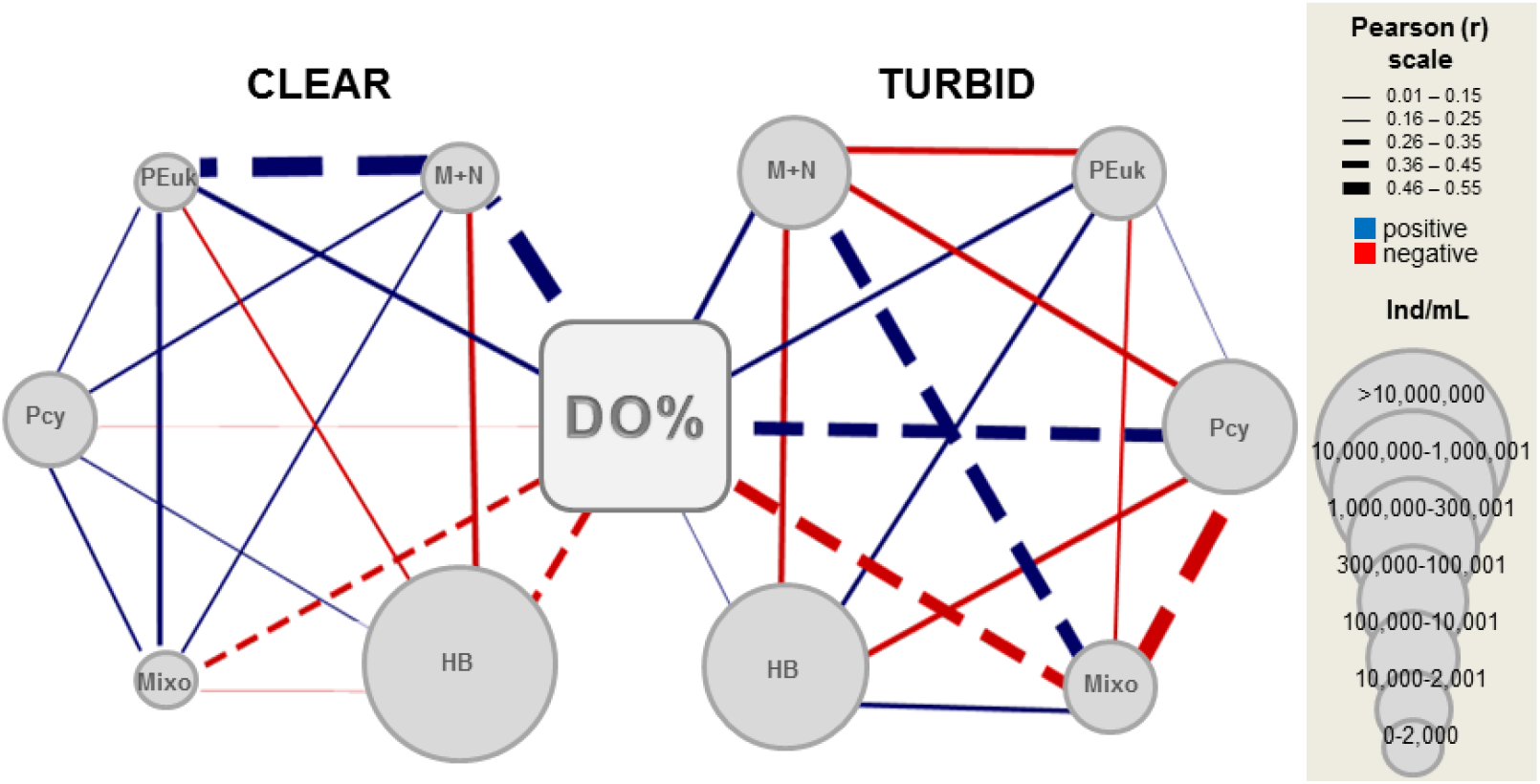
Correlation network among DO%, and micro + nanophytoplankton (M+N), picoeukaryotes (PEuk), picocyanobacteria (Pcy), mixotrophic algae (Mixo) and heterotrophic bacteria (HB) abundances under clear and turbid conditions at all times (T0, T1, T2 and TF). Circle size: mean abundance value (ind/mL) of each microbial community for the controls at T0. Dotted line: p<0.05. n = 72. Networks were built with Cytoscape^®^ v3.7.1 software.

## 4. Discussion

Our results strongly suggest that DO is sensitive for monitoring ecological changes driven by single and mixed herbicide formulations on freshwater systems. Despite the different modes of action of G and 2,4-D, their toxicity appeared to be primarily targeted at the algal and cyanobacteria fractions, probably because they share common pathways with plants (Couderchet & Vernet, 2003). Both herbicides, alone and/or in combination, can directly affect the autotrophic fraction of plankton in different ways, as already observed in microcosms (Lozano et al., 2018), leading to changes in DO dynamics in mesocosms.

Changes in DO levels in water may result not only from physical and chemical processes, but also from other environmental drivers of change acting on the biota, from individual to community scales. Contaminants such as herbicides may cause organisms to die or survive (Peck, 2011). The responses of the survivors depend on the rate of change and vary from biochemical buffering through gene expression and physiological flexibility at the individual scale, to changes in gene frequency, genetic drift and selection at the population scale. The latter are likely to occur relatively rapidly in microbial communities. Moreover, contaminants may induce stress responses in microbial communities, modifying their structure and function. These responses involve changes in respiration and photosynthesis rates, which affect the primary and secondary production of the ecosystem. Therefore, it is expected that herbicide inputs will increase the abundance of decomposers, thus decreasing DO concentration. An increased mortality of some autotrophic species may also cause a drop in DO concentration, as it contributes to additional decomposable matter and leads to a reduction in the photosynthetic rate. In contrast, physiological stimulation would increase the abundance of some autotrophic species, with the concomitant rise in photosynthetic rate and in DO concentration.

It is important to highlight that there are other aquatic microbial communities involved in the scenario of DO concentration changes driven by herbicides. Vera et al. (2010) found a strong impact of Roundup^®^ on periphyton, which is composed of autotrophic and heterotrophic fractions. On the other hand, Pizarro et al. (2015b) documented an increase in metaphyton (macroscopic mats of microscopic filamentous algae) biomass, following glyphosate addition. The impact of herbicides on freshwater systems may also include indirect effects on autotrophs. For example, Vera et al. (2012) reported that zooplankton preying on phytoplankton is affected by Glifosato Atanor^®^, a commercial glyphosate-based formulation commonly used in Argentina. Thus, variations in DO concentration reflect changes in the food web and in community structural and functional variables. Even if the overall picture of biological changes is incomplete, DO is a useful integrative variable that summarizes processes working in different directions.

The increase in DO at the higher concentration of G under the clear condition may be explained by the rise in the abundances of M+N phytoplankton and PEuk. In this sense, Pizarro et al. (2015a) showed a significant increase in PEuk in response to the input of Glifosato Atanor^®^. In the present work, the fact that no increase in PEuk abundance was induced at the low concentration of Roundup Max^®^ suggests a dose-response behavior. In contrast, in the turbid system, a similar pattern of increased DO would be related to an increase in Pcy abundance. Indeed, Pérez et al. (2007) observed that the addition of Roundup^®^ to earthen mesocosms triggered an increase in primary production related to a 40-fold rise in pico-cyanobacteria abundance. Pcy are autotrophic prokaryotes, with some species being tolerant to glyphosate (Pérez et al., 2007), regardless of water turbidity (Pizarro et al., 2015a), and capable of phosphonate degradation because they use it as phosphorus source (Ilikchyan et al. 2009, 2010).

There is very little information on the impact of AsiMax 50^®^ on freshwaters (Lozano et al., 2019) and only a few records concerning the effects of its active ingredient, 2,4-D, are available. The impact of 2,4-D on freshwater in general and on algal photosynthesis in particular is currently controversial, with some authors reporting stimulation (Kobraei & White, 1996; Boyle, 1980; Relyea, 2009) and others inhibition (Singh & Shrivastava, 2016; Zhu et al., 2016). In our study, the DO decrease in the turbid system at the high dose would be the result of the combination of a negative impact on one or more autotrophic populations and/or a positive impact on heterotrophic populations. The significant decline in M+N phytoplankton abundance observed at T2, that could have started before, and the increase in HB abundance at T0, were in accordance with the decrease in DO%. In contrast, neither DO% decrease nor variations in the phytoplankton fractions were observed in the clear system. After the addition of herbicides, changes in microbial communities occur relatively quickly as reported in previous studies (Pizarro et al., 2015a). As microbial communities respond quickly to impact factors (Peck 2011) they can recover quickly, depending on their own characteristics that can provide greater resilience (Oliver et al. 2015).

Our first hypothesis was related to the response of the two contrasting systems in terms of DO% dynamics under a similar contamination scenario while our second hypothesis was related to the similarity between the impact of Roundup Max^®^ and AsiMax 50^®^ on the daily DO%v dynamic. Glyphosate appeared to have an impact under both scenarios while 2,4-D had a stronger effect on DO dynamics only in the turbid system, hence, the first and the second hypothesis are rejected. Our results reveal that glyphosate, the most used herbicide worldwide, has significantly modified the dynamics of oxygen in water regardless of the type of system. This fact was already pointed out by Lozano et al. (2018), who showed that glyphosate had a higher effect than 2,4-D, either alone or in mixture, on the structure of phytoplankton and periphyton communities. Notwithstanding this, in our study G induced higher variations in the percentages of DO saturation and in the abundance of the phytoplankton in the clear system than in the turbid one. For example, in regard to abundance variation, the M+N phytoplankton showed a maximum mean value of 902% in the clear system, while the Pcy fraction reached a maximum of 58% in the turbid one under the GH treatment at T2. Although changes in microbial abundances due to glyphosate addition were higher in the clear than in the turbid system, values of DO%_v_ were not recovered at the end of the experimental period in the latter system. Changes in microbial abundances were lower in magnitude in the turbid than in the clear system, but the variation in the Pcy fraction in the last one would have had a higher influence, accounting for the DO% variations observed at TF with respect to control. The preference for phosphonates by the Pcy fraction has been already documented in different experimental approaches that were used to test glyphosate-based formulations in both clear and turbid systems (Pérez et al., 2007; Pizarro et al., 2015).

The network correlation revealed that the turbid system showed more correlations among the abundance of microbial fractions and DO%, suggesting that it is less resilient than the clear system for all herbicide treatments. A greater stability of the clear system under herbicide stress was also found by Lozano et al. (2019), who analyzed the impact of AsiMax 50^®^ on clear and turbid systems using a laboratory microcosms approach. The fact that in the clear system DO% was highly correlated with M+N phytoplankton abundances, while in the turbid one it was correlated with Pcy abundances may reflect different community responses to the herbicide (Cottingham et al., 1998) and supports the fifth hypothesis of this work. These community-related responses may depend on their structure and on the ability of each component to take advantage of the additional phosphorus provided by G. In terms of competition, it is probable that the M+N phytoplankton fraction was favored by the supply of nutrients under clear conditions, while the Pcy fraction was favored by light and nutrients under turbid conditions. The latter statement should be taken with caution considering that there are many other factors that are involved in the DO dynamics, such as the water chemistry and the fate of each contaminant in the environment.

The third hypothesis postulated that the DO%_v_ pattern followed an additive model in the presence of both herbicides. It was supported by all the results presented here, except for the synergistic effect observed at the high mixture concentration in the turbid system during the course of the experiment. A synergistic effect of a mixture of G and 2,4-D on phytoplankton structure has also been described in the context of freshwater communities by Lozano et al. (2018).

The fourth hypothesis that the impact of herbicides is stronger at high herbicide doses was supported by our results because the greatest variations in both DO% and abundance were observed in the treatments at high concentration. A dose-dependent impact of these herbicides on structural features of microbial communities has also been found previously (Lozano et al., 2018, 2019).

In our outdoor study, all the units in each experiment were subjected to the same climatic conditions (i.e., temperature, atmospheric pressure, rainfall and wind). Therefore, it can be assumed that DO% variations in water were exclusively determined by the impact of herbicide formulations on the structure of microbial communities and functional properties of the systems. To assess the effect of active principles and adjuvants present in commercial formulations on DO% variations, the present work focused on autotrophic microbial fractions and heterotrophic bacteria. However, other microbial fractions such as colorless flagellates, protozoa and rotifers should also be considered to draw more accurate conclusions (Vera et al., 2012). Certainly, biological interactions (e.g. competition, grazing) are known to play a relevant indirect role in DO dynamics.

Taking into account that most studies assessing the effect of binary mixtures of herbicides have dealt only with monospecific experiments (Backhaus et al., 2004), our work using two contrasting freshwater conditions at the ecosystem level is expected to provide more realistic results. The synergistic decrease of dissolved oxygen in the water due to the input of a mixture of herbicides raises concerns about environmental risks.

DO fluctuations in aquatic ecosystems may have several ecological consequences, including alterations in food web structure. Changes in DO have been assumed to affect fish behavior (difference of 1 mg/L) (Davis, 1975) and microbial stratification (Yu et al., 2014). On this basis, the magnitude of DO fluctuations measured in our study, with maximum values of 2.55 mg/L and 3.57 mg/L for clear and turbid scenarios, respectively, are expected to have relevant effects on for freshwater ecosystems.

Knauer and Hommen (2012) discussed the importance of functional vs. structural variables in assessing the collateral damage of herbicides in mesocosm studies. They concluded that functional variables generally show less experimental variability between replicates and provide a better integration of ecosystem function than do structural ones. Our results suggest that a simple metrics of DO may be sufficiently flexible to monitor the effects of herbicide mixtures on aquatic ecosystems regardless of their trophic status. This is especially true for outdoor mesocosm studies with all EUs having almost identical physical properties (Caquet et al., 2001). It is important to underline that a daily follow-up of DO concentration (e.g., with submersible data loggers) provides additional useful information. In freshwaters, DO monitoring is an effective tool to prevent environmental contamination, to evaluate the recovery of heavily polluted ecosystems and to analyze the potential impact of herbicides on the source-sink balance (Seguin et al., 2002). An assessment of the impact of agrochemical residues on water bodies is crucial for environmental protection, thus preserving the goods and services they provide to people. In addition, DO measures can give early and rapid warning of changes in the structure and abundance of communities from perturbed lentic freshwater systems.

Even though the variations in DO recorded as consequence of herbicide inputs may seem small, we believe that they are strong evidences of what would be happening in natural systems. Shallow lakes and rivers immersed in an agricultural matrix, such as that of the Pampa plain in Argentina, which are continually subjected to the arrival of herbicides, would have their DO dynamics also affected by these agents. Under a global warming scenario of increasing average temperatures and more frequent and longer droughts, water pollution by herbicides may contribute to anoxia, thus threatening water quality and aquatic life.

## Acknowledgments

This work was supported by PICT 2014.1586, UBACyT 20020130100248BA and PIP 11220130100399. The authors declare that they have no conflict of interest. This article does not contain any studies with human participants or animals performed by any of the authors.

